# Genetic Diversity, Population Structure and Species Delimitation of *Trialeurodes vaporariorum* (greenhouse whitefly)

**DOI:** 10.1101/067769

**Authors:** J. M. Wainaina, P. De Barro, L. Kubatko, M. A. Kehoe, J. Harvey, D. Karanja, L. M. Boykin

## Abstract

Genetic diversity within *Trialeurodes vaporariorum* (Westwood, 1856) remains largely unexplored, particularly within regions of Sub-Saharan Africa. In this study, *T. vaporariorum* samples were obtained from three locations in Kenya: Katumani, Kiambu and Kajiado counties. DNA extraction, PCR and Sanger sequencing were carried out on ~750 bp fragment of the mitochondria cytochrome c oxidase I (COI) gene from individual whiteflies. In addition, global populations were assessed and 19 haplotypes were identified, with three main haplotypes (Hp_19, Hp_10, Hp_011) circulating within Kenya. Measures of genetic diversity among *T. vaporariorum* populations resulted in haplotype diversity of 0.411, nucleotide diversity 0.00096, and Tajima’s D -0. 30315, (P>0.10). Analysis of population structure across global sequences using Structurama indicated one population globally, with posterior probability of 0.72. Bayesian and maximum likelihood phylogenetic analysis gave support for two clades (Clade I = an admixed global population and Clade II = subset of Kenyan and 1 Greek sequence). Species delimitation between the two clades was assessed by four parameters; posterior probability, Kimura’s two parameter (K2P), Rodrigo’s P (Randomly distinct) and Rosenberg’s reciprocal monophyly (P(AB). The two clades within the phylogenetic tree showed evidence of distinctness based on; Kimura two parameters (K2P) (p = -1.21E-01), Rodrigo’s P (RD) (p =0.05) and Rosenberg’s P(AB) (p = 2.3E -13). Overall, low genetic diversity within the Kenyan samples is a likely indicator of recent population expansion and colonization with this region and plausible signs of species complex formation in Sub-Saharan Africa.

---

Whiteflies are important agricultural pests with a global distribution (Anderson *et al.,* 2004; Lapidot *et al.,* 2014). Their agricultural importance is associated with damage caused during feeding on the plant phloem and the production of honeydew (Colvin *et al.,* 2006; Prijović *et al.,* 2013). Honeydew serves both to reduce transpiration and as an inoculation point for saprophytic fungi which grows over the leaf surface reducing photosynthesis (Colvin *et al.,2006).* Among the most economically important whiteflies is the greenhouse whitefly, *Trialeurodes vaporariorum* (Westwood, 1856) which transmits a number of plant viruses of the genera *Crinivirus* and *Torradovirus* (Navas-Castillo *et al.,* 2011; Navas-Castillo *et al.,* 2014). Of these, the *Criniviruses,* Tomato chlorosis virus (ToCV) and Tomato infectious chlorosis virus (TICV) are of major economic importance in tomato production globally (Wisler 1998, Wintermantel *et al.,* 2009). In Africa, ToCV and TICV have been reported from Morocco and South Africa (EPPO, 2005) and more recently from within greenhouses in Sudan (Fiallo-Olivé, et al., 2011). The viruses are able to infect a wide host range beyond tomato (Fortes *et al.,* 2012). Production loss attributed to *T. vaporariorum* transmitted viruses such as the sweet potato chlorotic stunt virus are a major constraint on sweet potato production in Kenya, with uninfected sweet potatoes yielding 50% more production than infected plants (Ateka *et al.,* 2004; Miano *et al.,* 2008).

## Phylogeographical structure of *T. vaporariorum* in Sub-Saharan Africa (SSA)

Though *T. vaporarioum* is globally distributed, few studies have explored the phylogeographical structuring and genetic diversity of *T. vaporariorum* populations from Sub-Saharan Africa and in particular Kenya (Mound and Halsey 1978; Njaramb, 2000). The majority of phylogeographical studies of *T. vaporariorum* have been confined to temperate regions (Roopa *et al.,* 2012; Kapantaidaki et al., 2014; Prijović *et al.,* 2014). In most of these studies there was lack of distinct phylogeographical structuring that was further coupled with low genetic diversity.

Roopa *et al.,* (2012) used both mitochondrial COI (mtCOI) and nuclear markers to assess the genetic diversity and phylogeographical structuring of *T. vaporariorum* within populations in India. They reported no significant differences among *T. vaporarioum* populations. Kapantaidaki *et al.* (2014) supported these findings using a combination of mtCOI and secondary endosymbionts. In addition, Prijović *et al.* (2013) reported a similar lack of phylogeographical structuring across populations in Serbia and surrounding countries. However, some studies have reported population structuring in different regions, with several drivers implicated as influencing population structure. In particular, Ovčarenko *et al.* (2014) reported population structuring in Finland and attributed this to habitats, field versus greenhouses populations. Gao et al., (2014) reported population structuring across the different regions of China and suggested the primary driver was multiple introduction points into China. There have been no similar studies in southern Africa.

On the basis of these studies we posed the following question: Do *T. vaporarioum* samples from sub-Saharan Africa and in particular Kenya have distinct phylogeographical structuring compared to global populations?

## How the hypothesis will be tested

To help answer the question, we carried out field survey in 2014 across smallholder farms located in the Kenyan highlands, which produce primarily common bean *(Phaseolus vulgaris, var humilis Alef).* Using mtCOI we compared the genetic diversity found in Kenya with that previously observed for *T. vaporarioum* in other parts of the world using records stored in GenBank. We assessed the genetic diversity of the different *T. vaporarioum* global populations and explored signals of species complex formation; this was based on several species delimitation methods. We report for the first time the phylogeographical structuring of *T. vaporarioum* samples within Sub-Saharan Africa and in particular Kenya, compared to global *T. vaporarioum* samples.

## MATERIALS AND METHODS

### Sample Collection

Sample collection of whiteflies was carried out on private property, following informed consent from the landowners. Adult whiteflies were collected on common beans (*Phaseoulus vulgaris*) growing within smallholder farms in Kenya. Sampling was done in the short rain season of November 2014, across counties sub-counties in Katumani/Machakos, Kiambu and Kajiado (Fig 1). In each farm, common bean plants were examined for the presence of whiteflies across a ‘z’ transect. Whiteflies were collected using a mouth aspirator and aspirated directly into a falcon tube (BD, Bioscienes USA) with RNAlater (Sigma, USA). Whiteflies were then shipped to the University of Western Australia and stored at room temperature.

**Figure 1:**
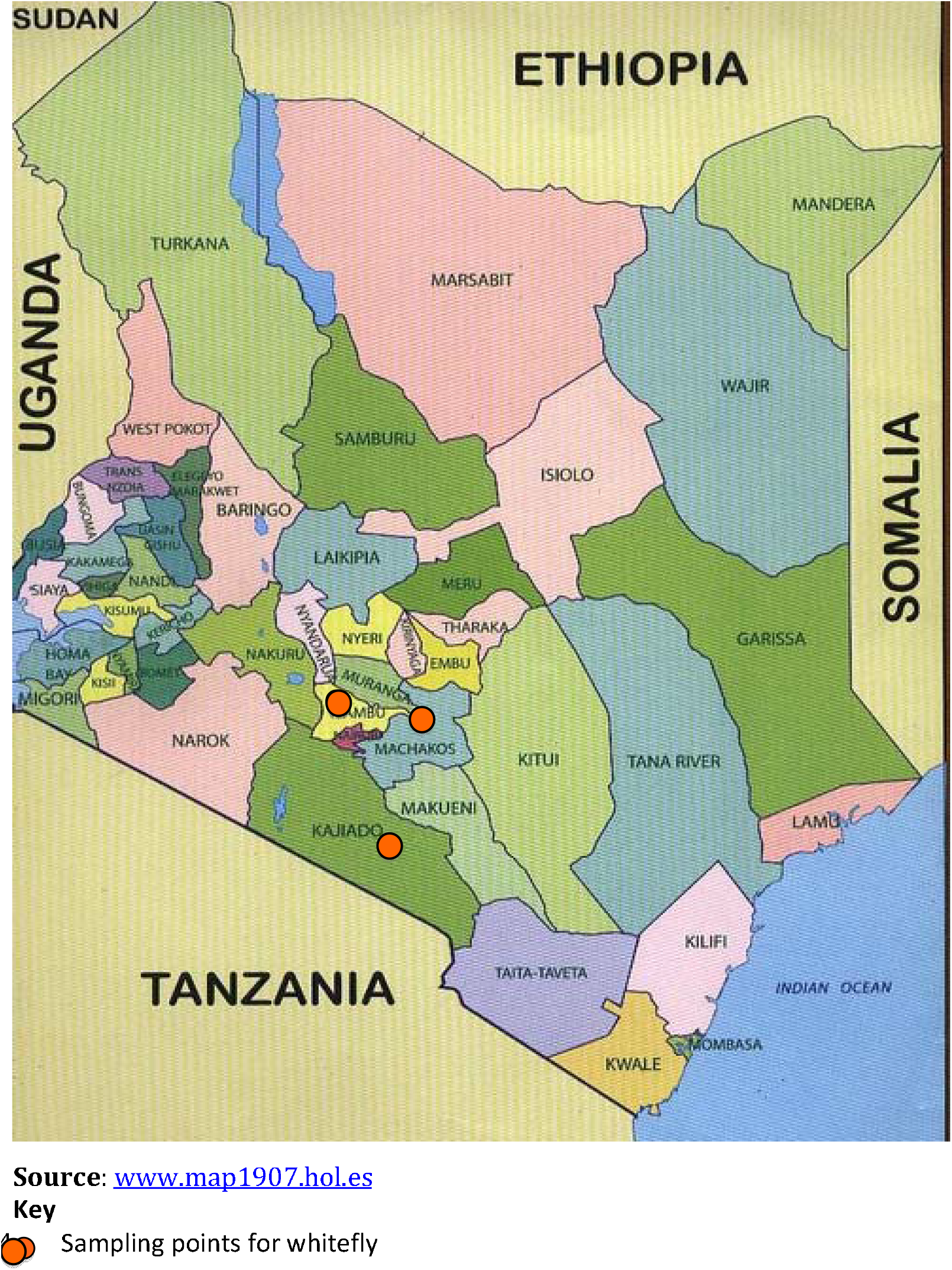
Map of Kenya with Sampling Points included in this study are highlighted

### Genomic DNA Isolation

Genomic DNA was extracted from individual whiteflies collected from each of the three sites using the Zymo Insect and Tissue extraction kit (Zymo, USA) as described by the manufacturer. Individual whiteflies were added into a ZR Bashing bead lysis tube containing 750 μl lysis solution. Bashing beads were then mounted on a Fast Prep®-24 MP bead beater and ground at 6 M/S for 40 seconds. Bashing beads were centrifuged at 10,000 × g for 1 minute. Subsequent steps were essentially as described by the manufacturer, with the only modification being the elution of genomic DNA with 30 μl of nuclease free water followed by a centrifugation step at 10,000 × g for 30 seconds. Genomic DNA was stored at -20°C until analysis was carried out.

### Polymerase Chain Reaction and Sequencing

Mitochondria COI *Trialeurodes vaporariorum* specific primers CO1-F: 5′-GCCTGGTTTTGGCATTA-3′ and the reverse primer CO1-R: 5′-GCTTATTTAGCACCCACTCTA-3′ were used to amplify ~752 bp of COI gene (Gao, et al., 2014). Polymerase chain reaction (PCR) was carried out in a total reaction mix of 20 μl with 5 ul of whitefly DNA as template, 10 ul Bulk AccuPower^(R)^ PCR Premix containing 1U of Top DNA polymerase, 250 μl of dNTP (dATP, dCTP, dGTP, dTTP), 10mM Tris-HCl (pH9.0), 30mM KCl, 1.5mM MgCl_2_, 1 μl of forward and reverse primers and 3 μl of nuclease free water. The PCR reaction was run on an eppendorf thermocycler (Eppendorf, Hamburg Germany) under the following conditions; Initial denaturation for 2 min at 94°C followed by 35 cycles of 94°C for 20 seconds, annealing for 30 seconds at 52°C, elongation for 1 minute at 72°C, final elongation for 30 min at 72°C. PCR amplicons were separated in a 2% agarose gel pre-stained with gel red (Biotium, CA, USA) and run in 0.5 X Tris Boric EDTA (TBE) electrophoresis buffer for 35 minutes at 100V and detected under UV light (Biorad, CA, USA). PCR products were purified using Qiaquick PCR purification kit (Qiagen Inc.) as described by the manufacturer. Purified PCR products were sent for bi-directional Sanger sequencing at Macrogen Korea.

### Sequence Alignment and GenBank Data Retrieval

A total of n=31 forward and reverse sequences of the 3’ end of mtCOI were obtained. Raw chromatograms were manually inspected using Geneious version 8 (http://www.eeneious.com, Kearse et al., 2012) across all the bases, with ambiguities corrected. Consensus sequences were generated for each whitefly, and used for subsequent analysis. Basic local alignment search (BLAST) was carried out to confirm that all sequences were *T. vaporariorum.* In addition, n = 228 *T. vaporariorum* COI sequences were retrieved from GenBank and combined with sequences from this study (Supplementary Table 1). The combined sequences were aligned using MAFFT in Geneious version 8 with default parameters (Kearse et al., 2012). Manual inspection, adjustment of gaps and trimming of overhangs was carried out.

### *Trialeurodes vaporariorum* Haplotype Distribution Across Geographical Locations

To better understand the distribution of *T. vaporariorum* haplotypes across geographical locations and reduce redundancy within COI sequences deposited in GenBank. *T. vaporarioum* COI sequences were categorised based on geographical locations considered to have similar climatic properties as illustrated within global ecoregion maps (Olson, et al., 2002) Representative sequences from each of the haplotypes and across the five geographical locations were selected creating a subset of 90 sequences for subsequent analysis (Table 2). Classification was as follows: location 1, - n = 40 sequences from Europe Asia, and North Africa; location 2, - n = 31 sequences from Sub-Saharan (Kenya) (this study), location 3 - n = 11 sequences from South America, Central America, and the Caribbean; location 4, India, China - n=7 sequences from India and China; and location 5 - n=1 sample sequences from North America (USA). Location 5 was excluded from all population analysis, since only one sequence could be retrieved from the GenBank.

**Table 1:** *Trialeurodes vaporariorum* across geographical location and host plant collected in this study.

| *Location | Coordinates | Host Plant | No of samples<br>sequenced | GenBank<br>Accession<br>Numbers |
| --- | --- | --- | --- | --- |
| **Kajiado** | -1.842073, 36.79186 | *Phaseolus vulgaris*<br>(Common Bean) | 2 | KX058204,<br>KX058211 |
| **Katumani** | -1.618715, 37.215651 | *Phaseolus vulgaris*<br>(Common Bean) | 4 | KX058202<br>KX058205<br>KX058207<br>KX058214 |
| **Kiambu** | -1.280825, 36.652756 | *Phaseolus vulgaris*<br>(Common Bean) | 25 | KX058201-KX058203<br>KX058206<br>KX058208-KX058209<br>KX058210<br>KX058212<br>KX058213<br>KX058215-KX058231 |

\*Location is the county or sub county in Kenya where whiteflies were collected in

**Table 2:** *Trialeurodes vaporariorum* haplotypes from across geographical locations determined using GenAlex 6.502

| Haplotypes | Location 1 | Location 2 | Location 3 | Location 4 | Location 5 | N | *GenBank<br>Accession<br>Number |
| --- | --- | --- | --- | --- | --- | --- | --- |
| Hp-1 | - | - | 1 | - | - | 1 | JF682887 |
| Hp-2 | 2 | - | - | - | - | 2 | KC843068 |
| Hp-3 | 6 | - | - | - | - | 6 | KC843065 |
| Hp-4 | 2 | - | - | - | - | 2 | KC843067 |
| Hp-5 | - | - | 1 | - | - | 1 | JF682883 |
| Hp-6 | - | - | 1 | - | - | 1 | JF682882 |
| Hp-7 | - | - | 1 | - | - | 1 | JF682881 |
| Hp-8 | - | - | 1 | - | - | 1 | JF682885 |
| Hp-9 | 4 | 23 | 2 | - | - | 29 | LN614547 |
| Hp-10 | 1 | 7 | - | - | - | 8 | KX058224 |
| Hp-11 | - | 1 | - | - | - | 1 | KX058231 |
| Hp-12 | - | - | 1 | - | - | 1 | JF682888 |
| Hp-13 | 1 | - | - | - | - | 1 | KJ475455 |
| Hp-14 | 1 | - | - | - | - | 1 | KJ475454 |
| Hp-15 | 2 | - | - | - | - | 2 | KF921192 |
| Hp-16 | - | - | - | 2 | - | 2 | JX841223 |
| Hp-17 | 1 | - | - | - | - | 1 | HE863766 |
| Hp-18 | 20 | - | 2 | 4 | 1 | 27 | KC113579 |
| Hp-19 | - | - | 1 | 1 | - | 2 | JQ995233 |

\*Representative GenBank Accession number provided for each haplotype.
\*N= Total Number of Haplotypes across the five world *Trialeurodes vaporariorum* populations615 **Geographical Location 1:** Europe Asia, and North Africa616 **Geographical Location 2:** Sub Saharan (Kenya, this study)617 **Geographical Location 3:** South America, Central America, and the Carribean618 **Geographical Location 4:** India and China619 **Geographical Location 5:** North America (USA)

### Inter-Population Genetic Diversity

The number of haplotypes within *T. vaporariorum* sequences was determined using GenAlex (Peakall & Smouse, 2012). mtCOI variation across the sequences was assessed using DNAsp v 5.0 (Librado & Rozas, 2009). Default program parameters were used during both the analyses.

### Haplotypes Networks Based on MtCOI Sequences of *Trialeurodes vaporariorum*

Association between haplotypes across the four global geographical locations were determined using median joining algorithms implemented in NETWORK version 5.0 (Bandelt *et al.,* 1999) with two outgroups *T. ricin and T. lauri* included under default parameters.

### Phylogenetic Analysis

Selection of the optimal model for phylogenetic analysis was determined using jModeltest 2 (Darriba *et al.,* 2012), using 11 substitution schemes. The optimal model selected based on the Akaike information criterion was (TPM2uf+G), while the optimal model in Bayesian information criterion was (HKY+I). The model HKY+I was selected since it considered variable base frequencies of one transition rate and one transversion rate. MrBayes was run on the Magnus Supercomputer at the Pawsey Supercomputer Centre at the University of Western Australia using 76 core/hours (274230s). Bayesian analysis was carried out for two independent runs using the command “lset nst=2 rates=gamma” for 50 million generations, with trees sampled every 1000 generations. In each of the runs, the first 25% (2,500) trees were discarded as burn in. Convergence of the runs was determined using Tracer vl.6.0 (Rambaut & Suchard, 2014). Maximum Likelihood (ML) phylogenetic trees with bootstraps and nucleotide model set at Ti ≠ Tv rate (2 ST) and were generated in PAUP* 4.0a147 (Swofford, 2003).

### Species Delimitation

Clades identified within the *T. vaporariorum* phylogeny were assessed as potential emerging species of *Trialeurodes vaporariorum* using the species delimitation plugin (Masters, Fan, & Ross, 2011) in Geneious version 8 (http://www.geneious.com, Kearse *et al.,* 2012). We used several species delimitation parameters on the mtCOI sequences; Kimura two parameter (K2P), Rodrigo’s (P(random distinct) posterior probabilities (liberal and strict) and Rosenberg’s P(AB) to evaluate either presence species delimitation within *T. vaporariorum* COI sequences (Table 5).

### Evaluation of *T. vaporariorum* Population Structure using Bayesian Analysis

To determine the population structure of *T. vaporariorum* mtCOI sequences across different geographical locations we analysed using Structurama 2.0 (Huelsenbeck, Andolfatto, & Huelsenbeck, 2011), which implements a Dirichlet process prior for the number of populations. The default parameters of Gamma (0.100, 10.000) were run, and the model without admixture was used. MCMC was run for one million generations and samples were collected every 1000 generations. Two independent runs were carried out to confirm that the MCMC algorithm converged to the same values (Table 4)

## Results

### Distribution of *T. vaporariorum* Haplotypes Across Global Populations

In this study, 31 *T. vaporariorum* sequences were generated and deposited in GenBank under accession numbers KX058201-KX058231 (Table 1) and compared with 228 sequences obtained from GenBank (Supplementary Table 1). Analysis across the combined global dataset based on representative 90 sequences revealed 19 haplotypes (Table 2).

### Inter-Population Genetic Diversity

Comparison of the genetic variation within *T. vaporariorum* was based on 443 bp fragments of COI sequences across all the populations. A majority of the haplotypes were from geographical location 1 (Europe Asia, and North Africa), while geographical location 2 Sub Saharan (this study) had the fewest (n = 3) haplotypes (Table 2). Assessment of population expansion based on neutrality test showed low values for Tajima’s D statistics across all four geographical locations indicative of population contraction; location 1 (−1.58699, p > 0.10), location 2 (−0.30315, p > 0. 10), location 3 (0.00351, p > 0.05) and location 4 (−1.23716, p>0.10) (Table 3), however none of the values were found to be statistically significant (Table 3).

**Table 3:** Variation of mitochondria COI sequences from *Trialeurodes vaporariorum* across four geographical locations calculated using DNAsp v 5.0.

|  | Pop 1 | Pop 2 | Pop 3 | Pop 4 |
| --- | --- | --- | --- | --- |
| Number of Individuals | 40 | 31 | 10 | 7 |
| Number of Haplotypes (h) | 10 | 3 | 8 | 3 |
| Haplotype (gene) diversity (Hd) | 0.726 | 0.411 | 0.956 | 0.524 |
| Nucleotide diversity (per site) (Pi) | 0.00248 | 0.00096 | 0.00351 | 0.00129 |
| **Neutrality Tests** |  |  |  |  |
| Tajima's D | -1.58699 | -0.30315 | 0.00351 | -1.23716 |
|  | (P > 0.10) | (P > 0.10) | (P > 0.05) | (P > 0.10) |

\*Four of the five population were considered; Population 5 was dropped since only one *Trialeurodes vaporariorum* sequence present from the GenBank.

### Median Joining Networks Based on MtCOI Sequences of *Trialeurodes vaporariorum*

Assessments of genetic relationships among the 19 haplotypes identified (Table 2) was based on a median joining network of the COI sequences (Fig 2), showing two ancestral haplotypes highlighted in red, red/black colour, while the rest of the haplotypes diverged from them. Haplotypes 9 (HP_9) and 18 (HP_018) were the dominant haplotypes across all geographical locations. Within the location 2 (Kenya), haplotype 9, 10 and 11 (Hp_9, Hp_10, Hp_011) were present.

**Fig 2:**
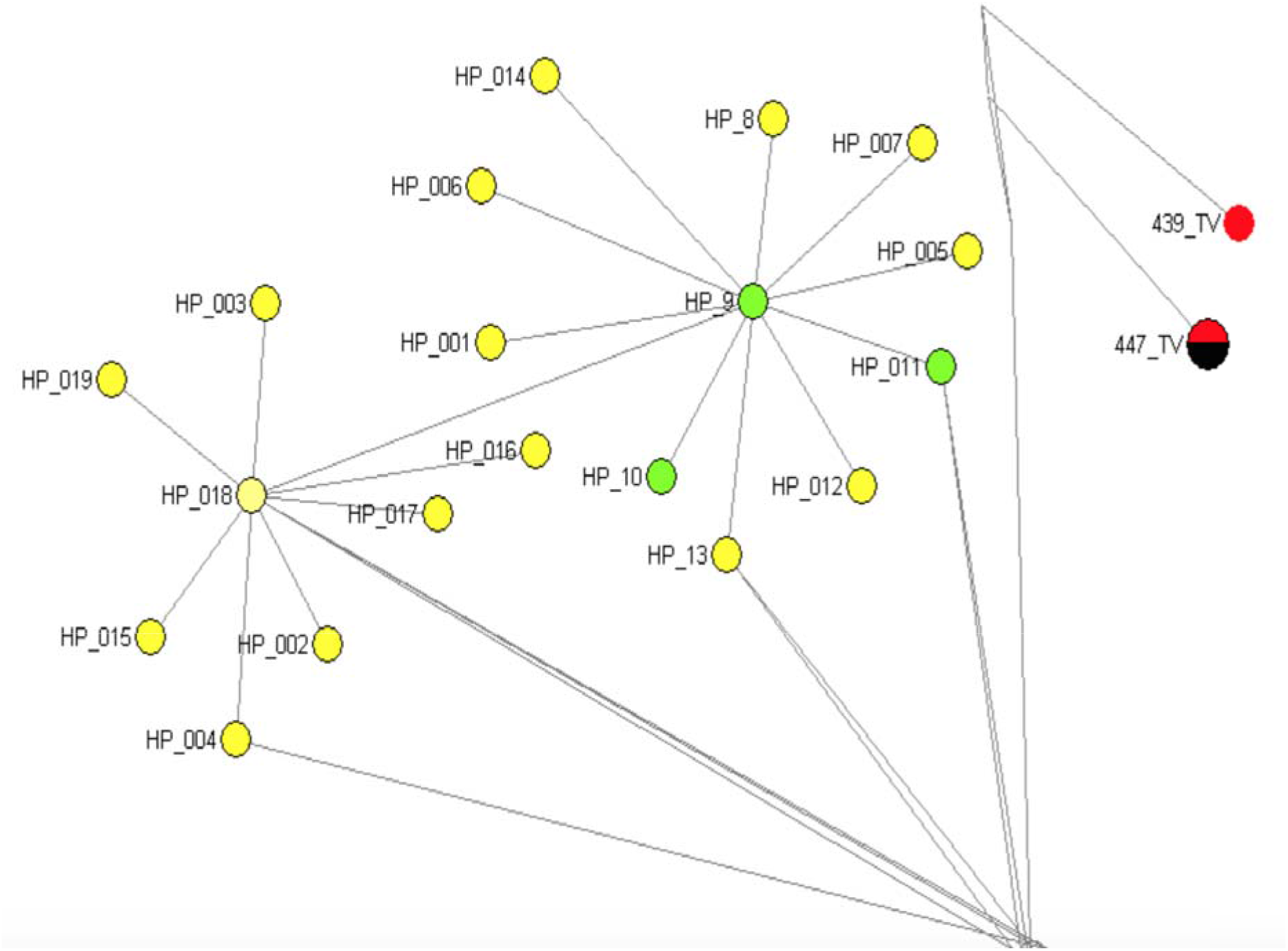
Median joining network for 19 MtCO1 haplotypes of *Trialeurodes vaporariorum* and 3 outgroup sequences (*Trialeurodes ricin, Trialeurodes lauri).* Median joining networks were generated using the program NETWORK version 5.0. Haplotypes in circulation in Kenya are highlighted in green while the ancestral haplotypes are in red/black.

### Phylogenetic analysis and species delimitation analysis of *Trialeurodes vaporariorum*

Phylogenetic relationships of *T. vaporariorum* across the geographical locations was assessed using Bayesian phylogenetic inference as implemented in MrBayes 3.2.2 (Huelsenbeck *et al.,* 2011) and maximum likelihood methods. Phylogenetic trees from both methods (Supplementary Fig. 1 and Supplementary Fig. 2) yielded two main clades, that are supported with 50% bootstrap support and >0.9 posterior probability respectively. Clade I was the largest clade, with a majority of *T. vaporariorum* including a subset of samples from geographical location 2 (SSA-Kenya). Clade II comprised of mainly *T. vaporariorum* individual sequences from geographical location 2 (SSA-Kenya) and one sequence from Greece. *T. ricin and T. lauri* were used as the outgroups for rooting the tree based on phylogenetic rooting methods described in (Kinene et al., 2016).

### Bayesian analysis of *Trialeurodes vaporariorum* Population Structure

Population structure of *T. vaporariorum* was evaluated using Structurama 2.0, with default parameter settings and no admixture. All of the *T. vaporariorum* sequences clustered as one population (K=1), supported by a posterior probability of 0.72 (Table 4), with posterior probability of 0.17 for K=2 populations. All other numbers of populations received posterior probability of less than 0.10.

**Table 4:** Posterior probabilities on the number of populations (K) within *Trialeurodes vaporariorum* for the COI sequences across geographical locations using Structurama 2.0.

| Number of Population<br>(K) | Posterior Probability |
| --- | --- |
| 1 | 0.72 |
| 2 | 0.17 |
| 3 | 0.07 |
| 4 | 0.02 |
| 5 | 0.01 |
| 6 | 0.0 |

### Species Delimitation

The two clades within the phylogenetic tree showed evidence of distinctness based on Kimura two parameters (K2P) (p = −1.21E–01), Rodrigo’s P (RD) (p =0.05) and Rosenberg’s P(AB) (p = 2.3E –13) (Table 5). Across the two clades all the sequences were monophyletic with genetic distance ranging between (0.2) and (0.7) (Table 5). The posterior probabilities of correct assignment of unknown species based on strict P ID (strict) and P ID (liberal) were (p >0.94) and (p >0.99) respectively (Table 5).

**Table 5:** Species Delimitation estimation using the species delimitation plugin in Geneious 8.1.8 on representative *Trialeurodes vaporariorum* across five global geographical locations

| Species | Closet Species | Monophyletic | Intra Dist | Inter Dist Closest (K2P(% difference)) | Intra/Inter | P ID(Strict) | P ID(Liberal) | Av (MRCA-tips) | P (Randomly distinct) | Clade Support | Rosenberg's P (AB) |
| --- | --- | --- | --- | --- | --- | --- | --- | --- | --- | --- | --- |
| 1 | 2 | Yes | -2.44E-03 | -1.21E-01 | 0.02 | 0.94 (0.83, 1.0) | 1.00 (0.94, 1.0) | -1.22E-03 | 0.05 | 83.56 | 2.3E-13 |
| 2 | 1 | Yes | -7.92E-03 | -1.21E-01 | 0.07 | 0.97 (0.92, 1.0) | 0.99 (0.97, 1.0) | -2.09E-02 | 0.05 |  | 2.3E-13 |

## DISCUSSION

In this study, we assessed the phylogeographical structuring and genetic diversity of *T. vaporariorum* within a previously underreported region (Sub-Saharan Africa). Phylogeographical structuring resulted in two main clades across representative global isolates with clade I (globally admixed) and II (Kenyan and one Greek sample). In contrast, genetic diversity across these global *T. vaporariorum* samples was low (Table 3). Species delimitation measures showed differences between *T. vaporariorum* isolates from Kenya compared to other geographical locations revealing plausible signs of species complex formation within a previously monophyletic group.

### Genetic Diversity of *T. vaporariorum* Across Global Samples

Genetic diversity of *T. vaporariorum* and haplotypes counts were based on a 443bp fragment of mtCOI sequence. A total of 19 *T. vaporariorum* haplotypes were present among all global sequences, with 3 haplotypes (Hp-9, Hp-10, Hp-11) present within *T. vaporariorum* from Kenya geographical location 2 (Table 2). Haplotype diversity ranged between 0.411 in population 2 to 0.956 in geographical location 3 (Table 3). Neutrality tests based on Tajima’s D statistic indicated low levels of genetic diversity across the global populations (Table 3). Our findings are comparable to previous genetic and population diversity studies of *T. vaporariorum,* indicating low genetic diversity (Roopa *et al.,* 2012, Shin *et al.,* 2013, Kapantaidaki *et al.,* 2014, Prijović *et al.,* 2014). Low genetic diversity within *T. vaporariorum* indicates recent population expansion and colonization into the regions. A possible explanation of the low genetic diversity of *T. vaporariorum* across global locations and in specifically from this study could be two-fold. Firstly, there has been an increase in global trade on horticultural products; mainly cut flowers from Kenya (Ulrich, 2014). This has resulted in increased importation of flower cuttings as propagation material from temperate ecoregions. Flower cuttings are potential carriers of *T. vaporariorum* because the eggs are invisible but readily hatch under ideal climatic conditions. This mode of *T. vaporariorum* movement has been implicated as one of the main modes of distribution of invasive pests globally (Grapputo *et al.,* 2005). Secondly, proliferation of polyhouses for horticulture production potentially provide appropriate microclimatic conditions sufficient for the emergence of *T. vaporariorum* and subsequent release into the open field. The importance of the polyhouse in building up *T. vaporariorum* populations and controlling gene flow have previously been reported in India, Northern and Southern Europe (Roopa *et al.,* 2012; Ovčarenko *et al,* 2014).

### Haplotype Networks

Evaluation of the relationship among the 19 *T. vaporariorum* haplotypes (Table 2) across five global geographical location based on the median joining network (Fig. 2) showed two ancestral nodes (Hp_09, Hp_018) (Fig. 2) that were present across all geographical locations. Within the SSA population three main haplotypes were in circulation (Hp_09, Hp_10, Hp_11) (Table 1) while Haplotype_11 was found as a distinct haplotype within this population. Our findings concur with previous reports on the number of haplotypes within *T. vaporariorum* (Prijović *et al.,* 2014). An overlap in haplotypes across the geographical location could indicate a single source of *T. vaporariorum* lineages that are easily adapted to different niche habitats (Kapantaidaki *et al.,* 2014). However distinct haplotypes and clades within location 2 (SSA-Kenya) could be indications of local adaptations of *T. vaporariorum.*

### Phylogeographical Structuring and Species Delimitation Genotypes

The evolutionary relationships among *the T. vaporariorum* sequences across geographical locations were reconstructed using both maximum likelihood and Bayesian phylogenetic analysis. Both phylogenetic trees resulted in two main clades (Supplementary Fig. 1 and Supplementary Fig. 2). Clade I was the largest clade composed of *T. vaporariorum* samples from all the geographical locations while clade II comprised of a subset of *T. vaporariorum* sequences from SSA (Kenya) and one Greek samples. This is contrary to previous phylogenetic relationships of *T. vaporariorum,* where a single clade was observed across *T. vaporariorum* populations collected (Roopa *et al.,* 2012; Prijović *et al.,* 2014; Kapantaidaki *et al.,* 2014). In this study, we report existence of at least two main clades within *T. vaporariorum* that are supported by bootstrap values of >60% and posterior probabilities above 0.90 (Supplementary Fig. 1 and Supplementary Fig. 2). Evaluation of the two clades as distinct species revealed signs of species complex within *T. vaporariorum* based on species delimitation measures (Table 5). Clade distinctness was supported by Kimura two parameters (K2P) (p = -1.21E-01), Rodrigo’s P (RD) (p =0.05) and Rosenberg’s P(AB) (p = 2.3E -13) (Table 5).

### Population structure based on Bayesian analysis

Population structure analysis of *T. vaporariorum* using Structurama based on a variable number of populations showed the strongest support for one population (posterior probability 0.72; Table 4). There was lack of distinct clustering from *T. vaporariorum* based on global geographical location (Table 5). Population structure results of one population were contrary to the phylogenetic tree showing the existence of two main groups within *T. vaporariorum* (Supplementary Fig. 1 and Supplementary Fig. 2). However, our finding concur with (Gao *et al.,* 2014) where they report lack of geographical structuring within the *T. vaporariorum* populations though several populations were reported. This further supports our hypothesis of a single source of *T. vaporariorum* introduced into Sub Saharan Africa.

Our findings highlight differences in the genetic diversity of the two whiteflies: *T. vaporariorum* and *B. tabaci,* which are both circulating in the regions sampled in Kenya. *B. tabaci* is recognized as a species complex of 34 species that are morphologically indistinguishable (Boykin *et al.,* 2007; De Barro & Ahmed, 2011; Boykin & De Barro, 2014). *B. tabaci* putative species exhibit differences in host range, phylogeographical clustering and potential to transmit viruses (Abdullahi *et al.,* 2003; Sseruwagi *et al.,* 2006, De Barro & Ahmed, 2011; Ashfaq *et al.,* 2014; Legg *et al.,* 2014). These differences are yet to be reported within *T. vaporariorum* and may be present based on distinct clustering observed and plausible species complex formation revealed in this study and would warrant further investigation.

Future studies should include a multilocus approach to provide for a more robust and stringent species delimitation calculation to authenticate the possible existence of a species complex formation within *T. vaporariorum.* However, it is probable that the existence of two clades are early indicators of impending divergence within *T. vaporariorum* potentially driven by phylogeographical factors as observed within *B. tabaci* and *T. vaporariorum* in Europe (Boykin *et al.,* 2013; Ovčarenko *et al.,* 2014). In addition, differences in the bacterial endosymbionts among *T. vaporariorum* haplotypes might induce genetic sweeps within the mitochondria and thus drive the diversification of *T. vaporariorum* (Kapantaidaki *et al.,* 2014). In addition, an expanded sampling from outside of Europe and throughout Kenya will greatly improve the understanding of this highly invasive pest.

## Conclusion

This study provides the evidence of phylogeographical structure of *T. vaporariorum* within Sub-Saharan Africa. In addition, the data presented here supports a putative species complex formation within *T. vaporariorum.* Our results provide preliminary findings for future research on the drivers for genetic population structuring and species complex formation within *T. vaporariorum* in nascent geographical regions.

**Fig 3.**
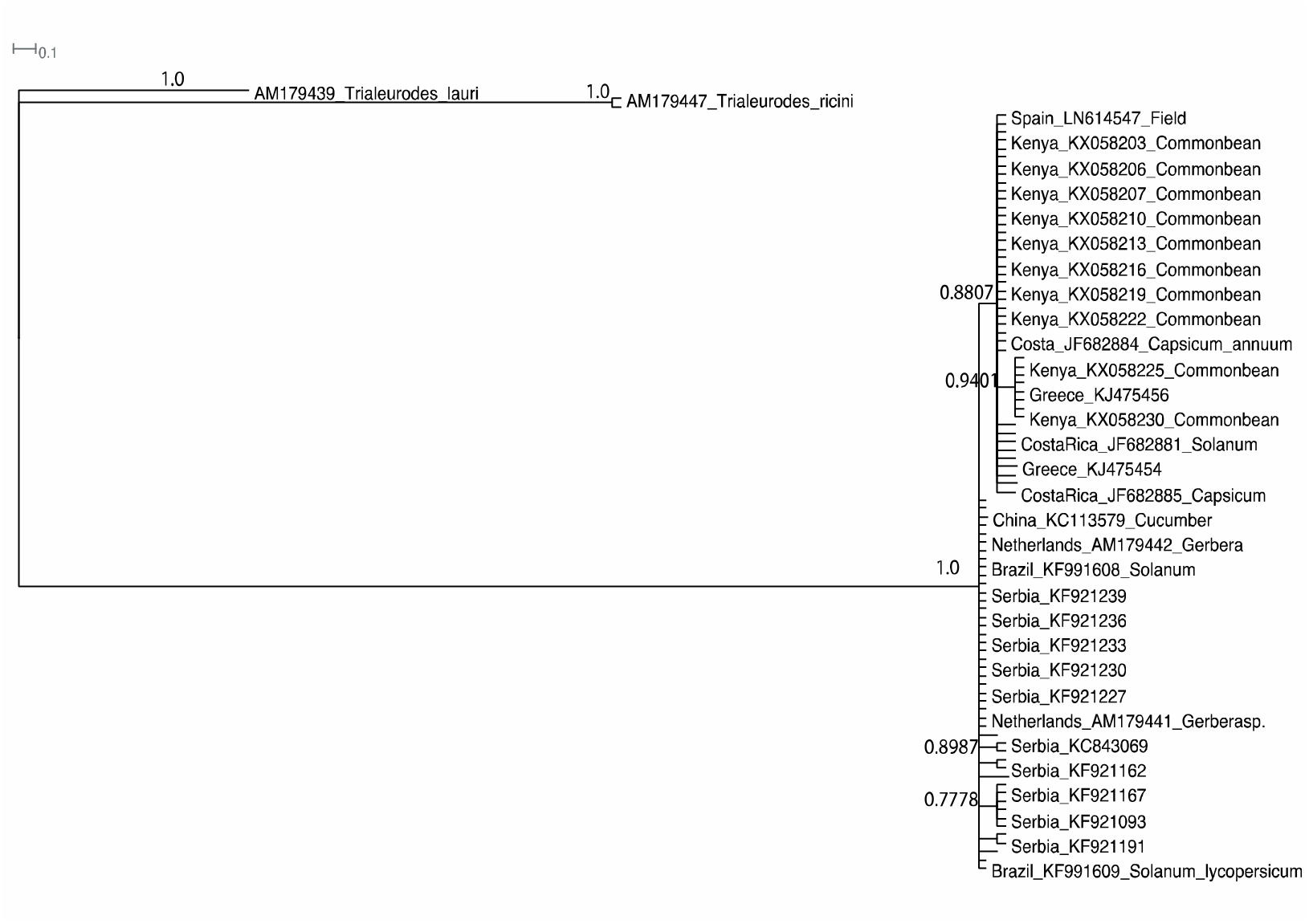
Bayesian phylogenetic relationships of representative *T. vaporariorum* and two outgroups (*T. ricin, T. lauri*) generated with MrBayes 3.2.2. Branches in supplemental figure 2 with zero branch length were collapsed.

## Acknowledgments

J.M.W is supported by an Australian Award scholarship by the Department of Foreign Affairs and Trade (DFAT), this work forms part of his PhD research. Fieldwork was supported by the kind consideration of Dr. John Carr through a BBSRC GRANT: BBSRC GRANT: BB/J011762/1 and the Bioscience eastern and central Africa (BecA-ILRI Hub). Laboratory work was performed at the Centre of Excellence Plant Energy Biology (PEB), The University of Western Australia. We thank Dr. Charles Kariuki Director Kenya Agricultural and Livestock Research Organization (Katumani) for facilitating sample collection in Eastern Kenya and insightful discussions. Supercomputer analysis was supported by resources provided by the Pawsey Supercomputing Centre with funding from the Australian Government and the Government of Western Australia. Research in the Boykin lab is provided by the Bill & Melinda Gates Foundation via a subcontract (B04265) from the National Research Institute, University of Greenwich, United Kingdom.

---

**Supplementary Table 1:**
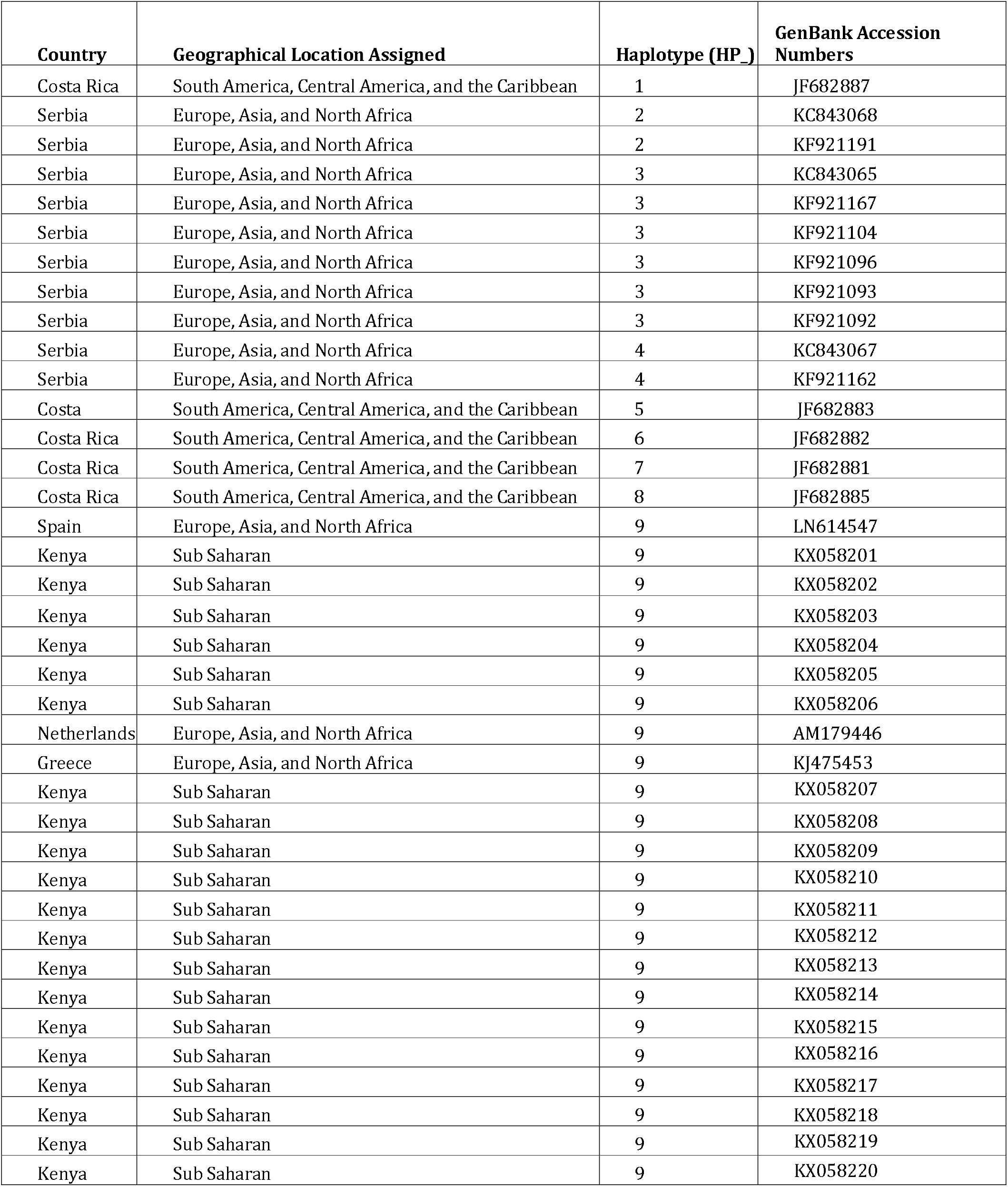
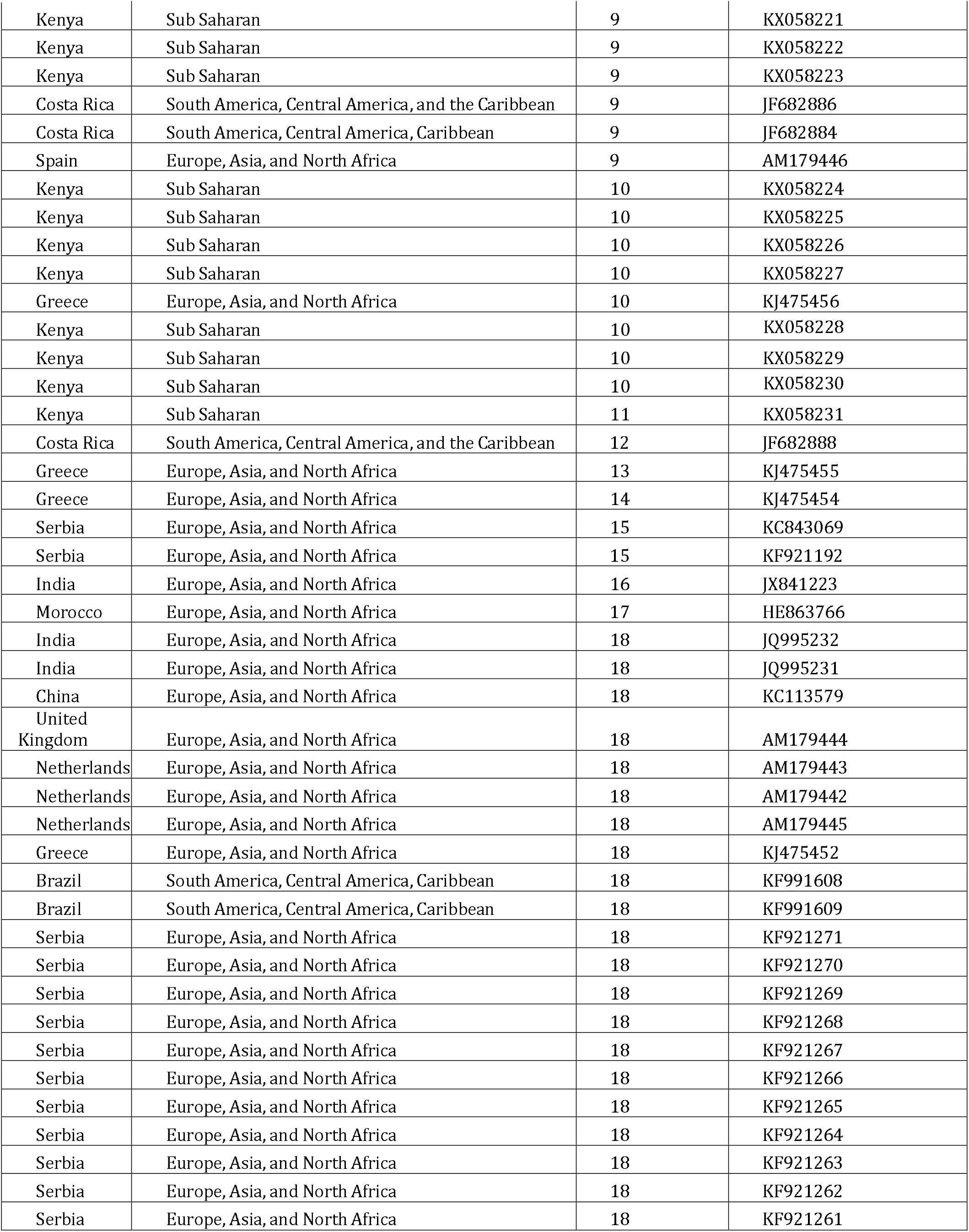
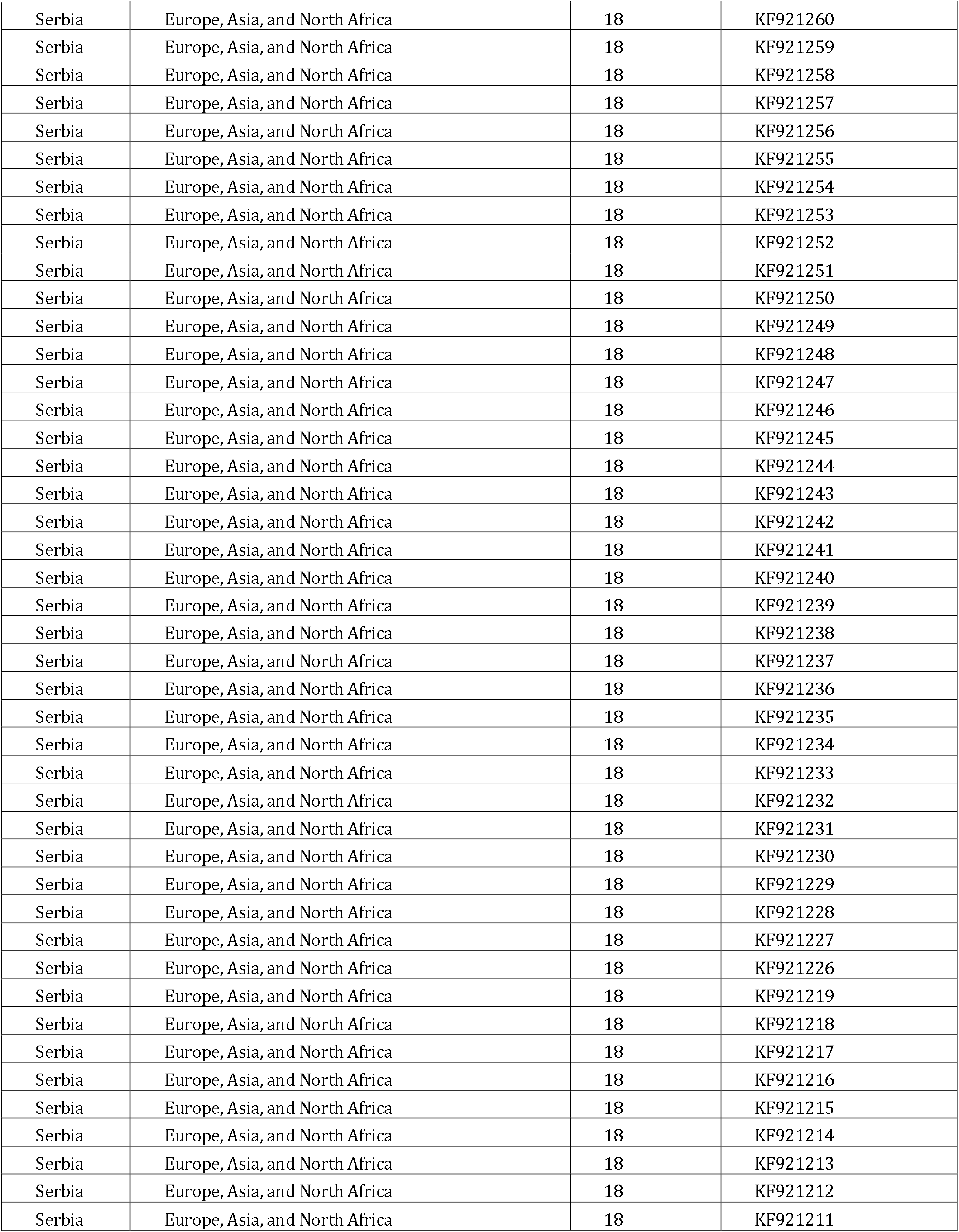
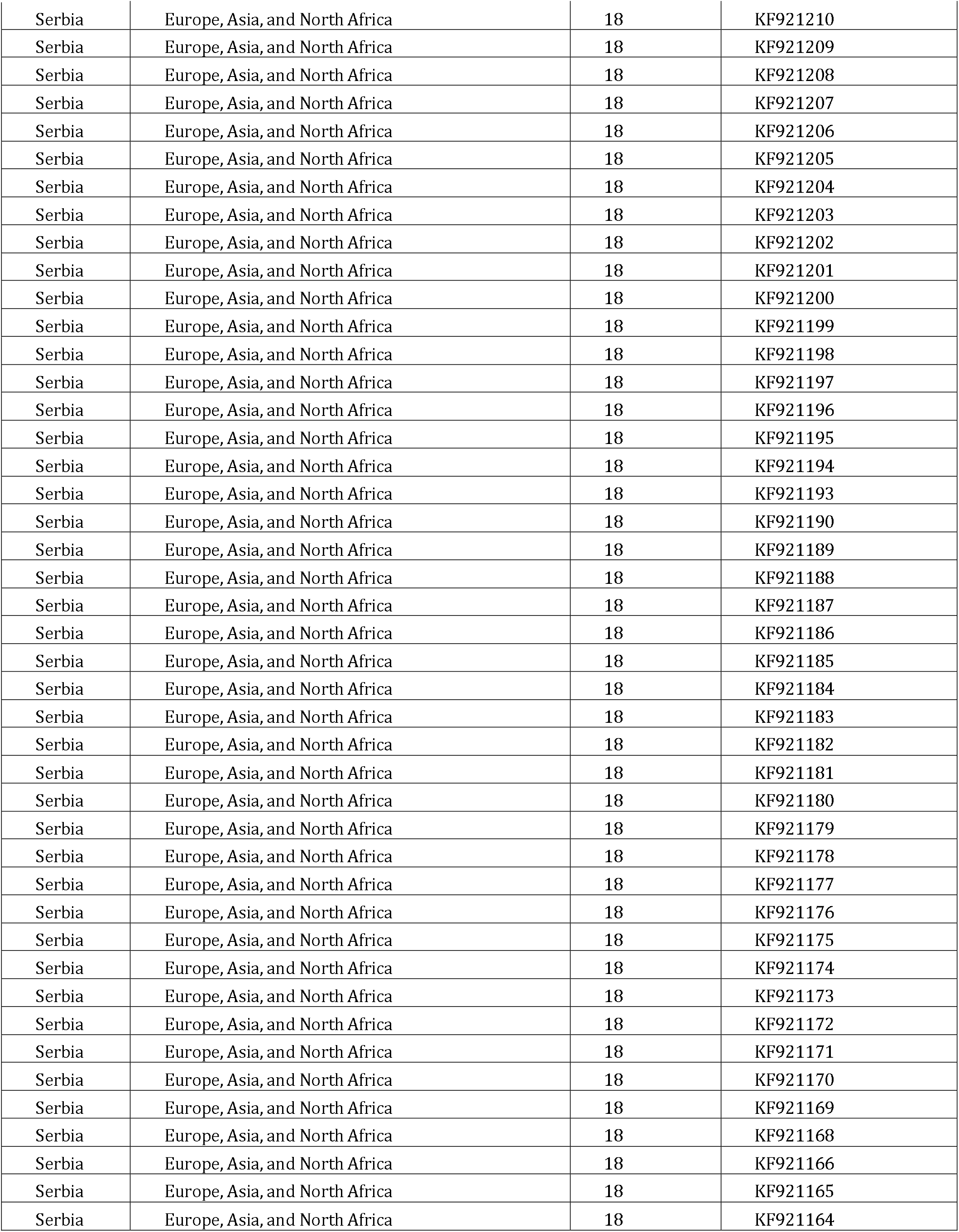
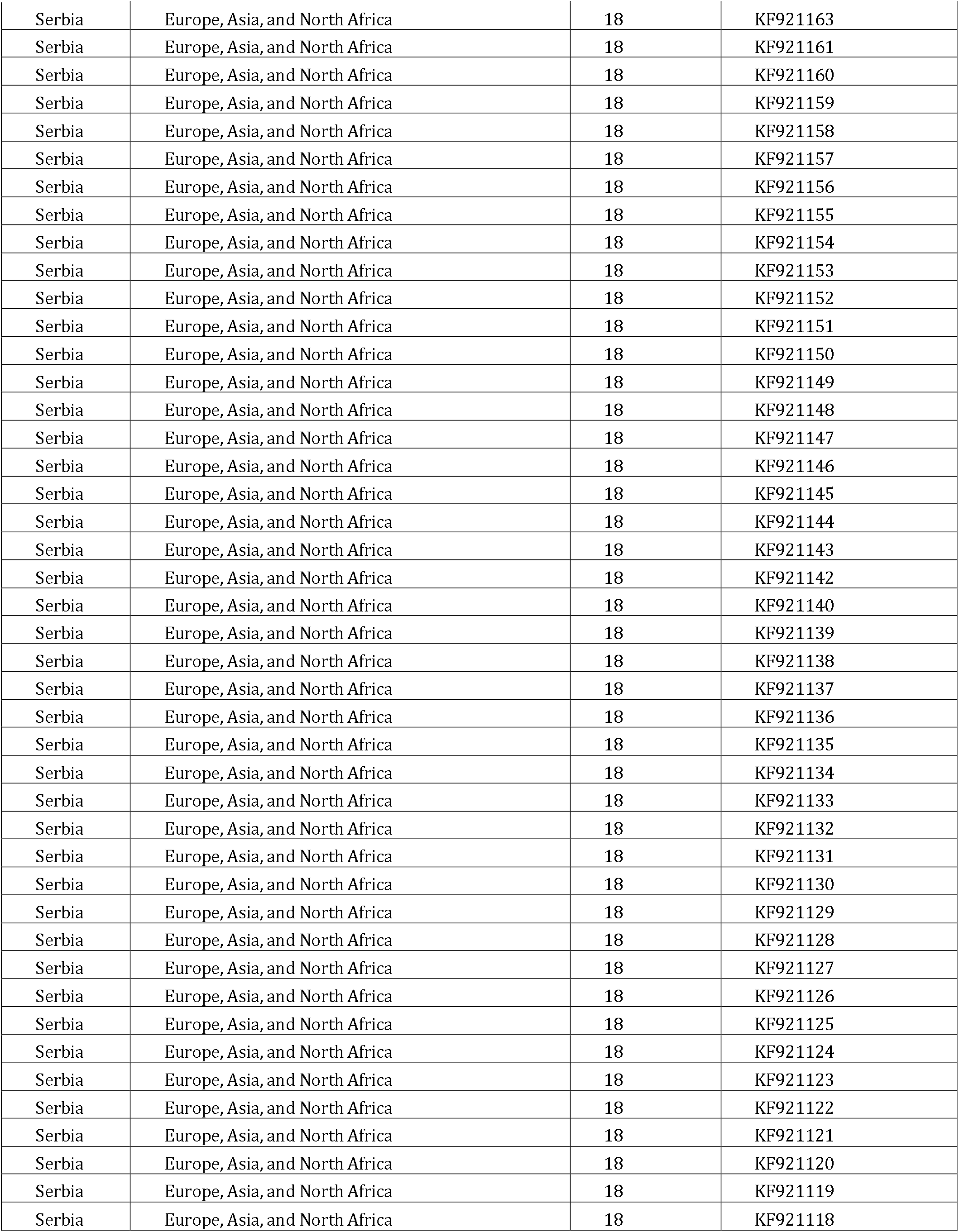
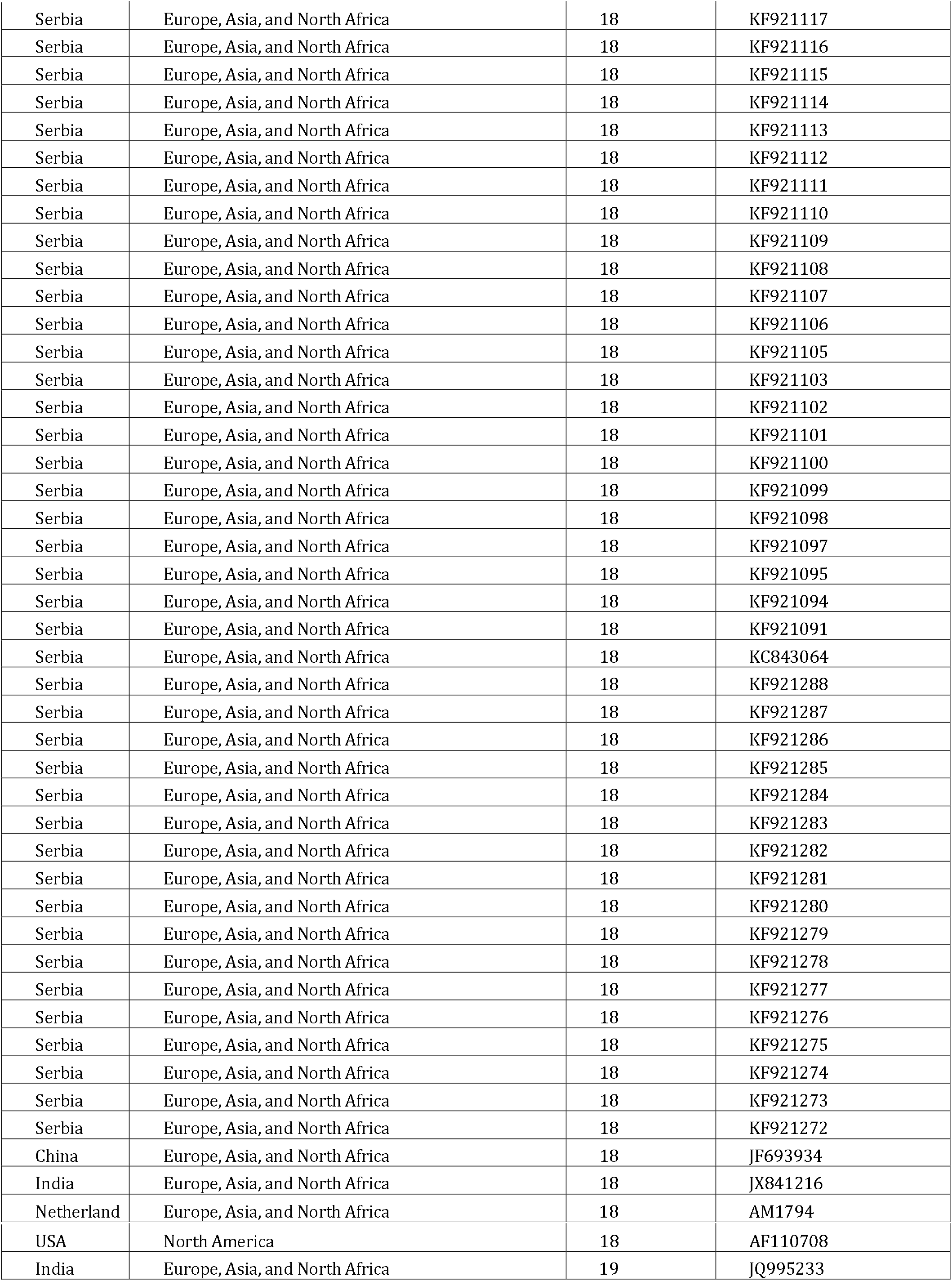
*Trialeurodes vaporariorum* COI sequence from GenBank and this study with respective haplotype and Geographical regions assigned

